# Identifying Key Determinants of SARS-CoV-2/ACE2 Tight Interaction

**DOI:** 10.1101/2020.07.13.199562

**Authors:** Van A. Ngo, Ramesh K. Jha

## Abstract

SARS-CoV-2 virus, the causative agent of Covid-19, has fired up a global pandemic. The virus interacts with the human receptor angiotensin-converting enzyme 2 (ACE2) for invasion via receptor binding domain (RBD) on its spike protein. To provide a deeper understanding of this interaction, we performed microsecond simulations of the RBD-ACE2 complex for SARS-CoV-2 and compared it with the closely related SARS-CoV discovered in 2003. We show residues in the RBD of SARS-CoV-2 that were mutated from SARS-CoV, collectively help make RBD anchor much stronger to the N-terminal part of ACE2 than the corresponding residues on RBD of SARS-CoV. This would result in reduced dissociation rate of SARS-CoV-2 from human receptor protein compared to SARS-CoV. This phenomenon was consistently observed in simulations beyond 500 ns and was reproducible across different force fields. Altogether, our study shed light on the key residues and their dynamics at the virus spike and human receptor binding interface and advance our knowledge for the development of diagnostics and therapeutics to combat the pandemic efficiently.

## INTRODUCTION

It is vital to understand the mechanisms of SARS-CoV-2 in comparison to other coronavirus.^1–8^ One strategy is to zoom in to the molecular levels of how SARS-CoV-2 invades human cells. The first step of the invasion is to have its Spike proteins bind to human cells such as lung cells, whose surface expresses a lot of ACE2. The RBD of SARS-CoV-2^2^ (dubbed RBD2 in this study) was identified to bind to ACE2 with an interface similar to the RBD of SARS-CoV (dubbed RBD1 hereafter) from 2003.^9^ It was determined that RBD2 binds stronger to ACE2 than RBD1.^4,10,11^ There are multiple mutations that are found in RBD2 that can explain why SARS-CoV-2 is more infectious than SARS-CoV. Specifically, residues G482/V483/E484/G485/F486/Q493/L455/N501 (based on RBD2 numbering) constituting a receptor binding motif (RBM) may cause a significant structural difference when comparing the crystal structures of RBD2 and RBD1.^2^ Free-energy calculations were also done for the mutations at these residues to compare how they contribute to overall binding affinities via a coarse grain model.^12^ Using homology models for comparing the structures of Bat-CoV, SARS-CoV, and SARS-CoV-2, Ortega et. al.^5^ proposed that the two loops of RBD2 might help enhancing its interactions with ACE2 with an improvement of 1.6 kcal/mol in the binding energy compared to RBD1. Amin et al.^13^ showed this increase in the binding affinity may be attributed to the electrostatic interactions enhanced by the mutations appearing in RBD2. Most of these studies relied on short simulations, thus not revealing the dynamics of the interfaces. To aid the research community, the Shaw group^7^ performed 10-75 μs molecular dynamics (MD) simulations of both RBD1 and RBD2 and how they bind to ACE2, and has granted a free access to many critical simulation data. However, how these residues change the conformations at the RBD2-ACE2 interface in comparison to the RBD1-ACE2 interface remains to be elucidated. Even though free-energy calculations^12^ reveal relatively important levels of these residues at RBD2-ACE2 interface, how they dynamically co-operate to enhance the binding of RBD2 and how RBD2 may dynamically dissociate from ACE2 have not yet been explored.

In this study, we demonstrated how the key residues F486/ N487/Y489/A475, some of which were previously identified as important, play a more critical and collective role at RBD2-ACE2 interface than other residues (for example, Q493/L455/N501) and the corresponding residues L486/N487/Y48/P475 at RBD1-ACE2 interface (**Figure 1a**). We cross-examined these residues in two popular force fields (FFs), CHARMM36^14^ and AMBER FF.^15^ Understanding how these groups of residues dynamically and collectively respond to an applied “force” at RBD2-ACE2 interface may lead to an effective therapeutic design to specifically target these residues for battling the invasion of SARS-CoV-2 to the human host cells. To show such a collective response, we performed Umbrella Sampling (US) simulations of 4.8 μs in total per complex to compute the potential of mean forces (PMFs) for separating the RBDs from ACE2. Combining the results from long equilibration simulations and US simulations, we demonstrated that the dynamics of the residues at RBD1-ACE2 and RBD2-ACE2 interfaces are quantitatively different, that is, the loop, where F486/N487/Y489/A475 are located in RBD2, anchors strongly on ACE2, but the corresponding loop in RBD1 may lose its grip on ACE2.

**Figure 1.**
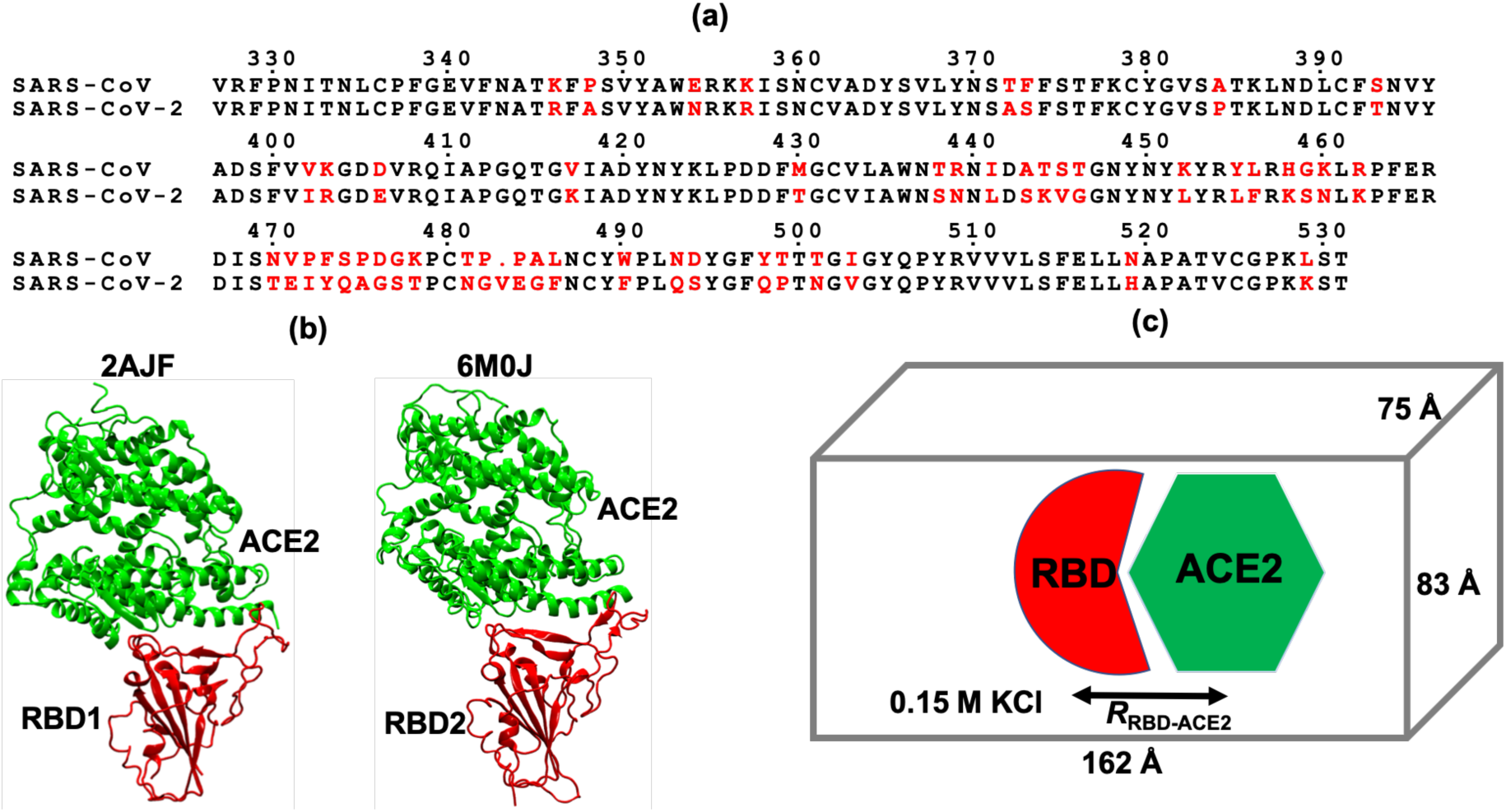
(a) Sequence alignment of receptor binding domains (RBD) of SARS-CoV and SARS-CoV-2. The residues highlighted in red are mutations found in the RBDs. Any residue numbers referred in the text are positions in this sequence alignment. This alignment is consistent with the alignment reported in Ref.^4^ (b) X-ray structures of RBD1 of SARS-CoV (PDB:2AJF)^9^ and RBD2 of SARS-CoV-2 (PDB:6M0J),^2^ respectively, bound to human receptor angiotensin-converting enzyme 2 (ACE2). (c) MD simulation setup for RBD1-ACE2 and RBD2-ACE2 complexes.

## METHODS

For the MD simulations, we used the crystal structure (PDB:2AJF)^9^ for RBD1-ACE2 complex, and the crystal structure (PDB:6M0J)^10^ for RBD2-ACE2 complex (**Figure 1b**). Note that the residue numbers in **Figure 1a** are consistent with the alignment in Ref.^2,4,6^. For simplicity and consistency, we refer to the same residue numbers for both complexes. The residue indices in the PDB:2AJF can be converted to the residue numbers shown in **Figure 1a** by adding 13 to indices less than 472 or by adding 14 to the indices otherwise since RBD2 showed an insertion of a residue at position 483. The missing loop (residues 375-381) in RBD1 was added by aligning RBD1 with RBD2; then the coordinates of the aligned loop (residues 389-394) was used to construct the coordinates for the missing loop in RBD1. We then used CHARMM 36 (C36)^14^ force field (FF) to model the complexes. When building the models, we did not include glycans in either of the complexes. The asparagine (N90) linked glycan (on ACE2), close to the interface, showed better electron density in RBD1-ACE2 structure and showed an extended form consisting of three sugar units (PDB: 2AJF). The closest residue on RBD1, T402, showed a distance of > 4 Å from the glycan and would contribute weakly to the interaction. In RBD2-ACE2, N90 linked glycan lacked electron density and consisted of only one sugar unit. Another structure of RBD2-ACE2 (PDB:6M17) also showed only two sugar units linked to N90. In both RBD2 complexes, the nearest amino acid to the glycan sugar T415 was not expected to contribute to the binding interaction. Due to this discrepancy and a low binding energy contribution from glycan, the complexes were prepared in the absence of them. We included the Zn^2+^ ion, which is coordinated by two histidine residues (H356 and H360) and a glutamate (E384) of ACE2. All histidine residues were modeled in the neutral and non-protonated state. We applied a patch of disulfide bond for all disulfide pairs as found in both RBDs and ACE2.^2,9^ The systems were then solvated with TIP3P water model.^16^ 0.15 M KCl was added to neutralize and mimic a physiological condition. The number of K^+^ and Cl^−^ were kept at 102 and 78, respectively, for simulations of both RBD1-ACE2 and RBD2-ACE2 complexes. The total number of atoms in these two solvated systems were approximately 102,000 with a dimension of 75×83×162 Å^3^ after minimization and 0.5 ns equilibration using NAMD 2.12.^17^ The minimization step was done with conjugate gradient energy minimization method.^18^ A cut-off distance of 12 Å and a switch distance of 10 Å was used to compute Lennard-Jones interactions. We enabled Particle Mesh Ewald summation method^19^ to compute electrostatic interactions with a grid-size of 1.0 Å. We used Langevin dynamics together with Langevin Piston to keep temperature around 310 K and pressure at 1 atm. A timestep = 1 fs was used for the short equilibration simulations.

To enable a larger integration timestep, we used PARMED^20^ to re-partition hydrogen atoms bonded to protein heavy atoms (exclude water molecules), while keeping the total mass unchanged.^21^ In this way, the dynamics of an entire protein may not noticeably change but the simulations can become more stable when using a timestep of 4 fs. We used AMBER version 16^22,23^ in order to run efficiently on the GPU nodes using the same C36 FF and simulation parameters used in NAMD 2.12. Specifically, we can pack four replicas per GPUs-node without a cost of slowing down each of the four simulations per node via AMBER, while the same setup for the simulations using NAMD-GPU can experience a significant slow-down (data not shown).

Since it has been shown that running multiple independent MD simulations in parallel can sample better than running a single long MD simulations,^24,25^ we created 8 replicas of each complex, which were quickly equilibrated using NAMD 2.12. Each replica was further equilibrated using AMBER 16 for about 600 ns to collect data, which were saved every 1 ns for analysis. We collected data for a total of approximately 5 μs per complex.

To compute a potential of mean force (PMF), which can be used to describe an effective interaction between the complexes as a function of the distance between the two centers of mass, we performed Umbrella Sampling (US) simulations (Figure 1b).^26–28^ Typically, US simulations are prepared in a set of independent simulation windows, each of which has an applied or biased harmonic potential, e.g., *U*_US_(*R*_*i*_,*R*) = *k*(*R*_*i*_ *– R*_RBD-ACE2_)^2^/2, where *k* = 10 kcal/mol-Å^2^, *R*_RBD-ACE2_ is the biased distance between the centers of mass of an RBD and of ACE2 (for simplicity, we used only Cα atoms to compute the centers of mass); and *R*_*i*_ is a restraint position. We found the smallest value of *R*_*i*_ from the distribution of *R*_RBD-ACE2_ from the above 4.3 μs equilibration simulation. For instance, in the case of RBD1-ACE2 complex, the smallest value of *R*_*i*_ is 43 Å; in the case of RBD2-ACE2 complex, the smallest value of *R*_*i*_ is 46 Å. Then, we chose a last window with *R*_*i*_ ∼ 70-75 Å to enable a complete separation between each RBD and ACE2. Since the harmonic potential induces a fluctuation of about 0.3-0.4 Å around each restraint position *R*_*i*_, we used an increment of 0.5 Å between every adjacent window (Supplementary **Figure S1**) to ensure proper overlaps between the distributions of *R*_RBD-ACE2_ in any two adjacent windows.^29^ Each window was run for 10-11 ns per replica, i.e., for 8 replicas, we collected approximately 80 ns per window. In total, we simulated for 60 windows with a total of 4.8 μs for each complex. We saved the values of *R*_RBD-ACE2_ every 1000 steps (=4 ps) and the biased configurations every 1 ns for each window. These values were then used to solve for a potential of mean force via Weighted Histogram Analysis Method.^27,28^

## RESULTS

### Key residues at RBD2-ACE2 interface

To rank the relative importance of each residue at the interfaces of RBD1 and RBD2 with ACE2, we computed the “bound” probability of each pair at the interfaces during the 5-μs equilibrium simulations. **Figure 2a-b** shows a relative ranking of pairs of residues at the interface: high probabilities indicate highly interactive pairs, while low probabilities indicate weakly interacting pairs. To the best of our knowledge, such a complete ranking has not been reported in previous studies, even though calculations of relative changes of interaction energies or free-energies were done for some of the pairs and mutations.^12,13^ **Figure 2a-b** shows that both interfaces have almost the same number of interacting pairs. However, if counting stable pairs that have probability larger than 60% (i.e., in close contact 60% of the total simulation time), there are only 22 ± 2 pairs in RBD1-ACE2 interface, while RBD2-ACE2 showed substantially higher, 35 ± 1 stable pairs at the interface. Among these stable pairs, the mutations from RBD1 to RBD2, namely, Y455L, L456F, Y498Q, T501N and L486F appear to noticeably enhance the probabilities of interaction with the residues of ACE2 with multiple pairs having probabilities close to 100%. The green brackets in **Figure 2b** indicate two groups of neighboring residue clusters that have a substantial increase in interactions in RBD2-ACE2 interface compared to the similar clusters in RBD1-ACE2 interface. This suggests that they are key residues that differentiate RBD2-ACE2 binding interface from RBD1-ACE2 binding interface.

**Figure 2.**
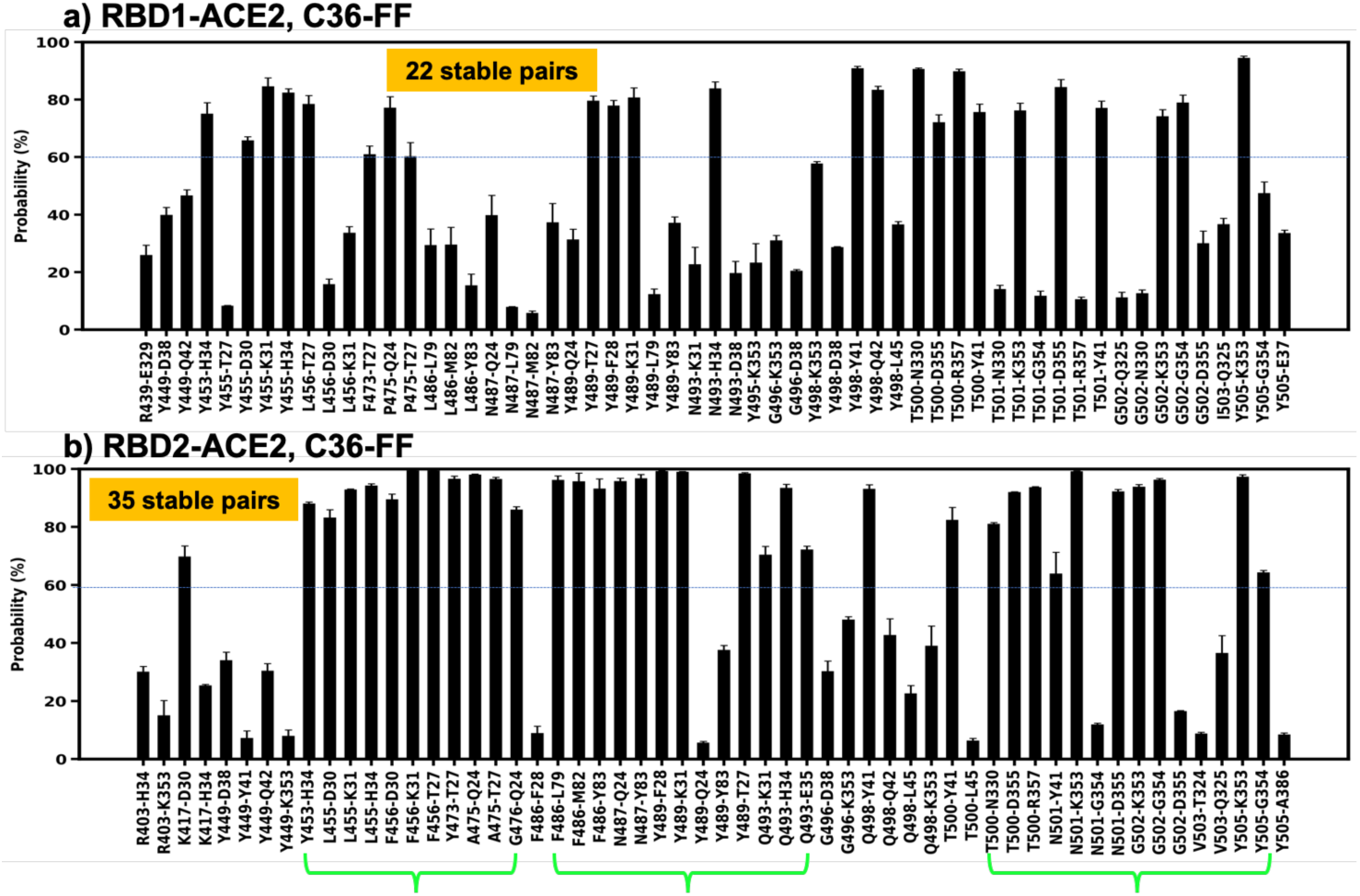
(a-b) Probability of pairs of residues found within 3.5 Å at the interfaces between RBD1 or RBD2 with ACE2 during simulations of 5 μs per system using C36 FF. A stable pair is defined to have a probability larger than 60% during the simulation times, i.e, forming a pair within 3.5 Å for more than 60% of a simulation trajectory. Green brackets indicate clusters of residues that are on average more stable in RBD2-ACE2 interface than the corresponding clusters in RBD1-ACE2 interface.

To probe whether the use of a different force field may change the outcomes of the key residues in the interfaces, we compared the probabilities with those obtained from the simulations using modified AMBER ff88SB FF^15^ performed by the Shaw group.^7^ In this AMBER-FF model, RBD1-ACE2 interface shows 29 ± 2 stable pairs, which are again substantially lesser than 39 ± 1 stable pairs in RBD2-ACE2 having bound probabilities larger than 60% (Supplementary **Figure S2**). As a result, the AMBER FF quantitatively yields more stable pairs that the C36 FF. Particularly, the AMBER FF predicts two salt-bridges (R403-E37 and K417-D30) in the RBD2-ACE2 complex, compared to only one salt-bridge when modeled by the C36 FF (K417-D30). However, the Y455L, L456F, Y498Q, T501N and L486F mutations (Supplementary **Figure S2**) are also found to enhance the same clusters of interactions in RBD2-ACE2 interface (**Figure 2b**). Hence, regardless of the FFs, the RBD2-ACE2 interface shows stronger inter-molecular interactions than the RBD1-ACE1 interface involving the same group of residue positions (**Figure 2b**). The clusters of the important residues are consistent with the finding^2^ that residues L455 and F486 and N501 of SARS-CoV-2 are perhaps the most critical residues that increase the interactions between RBD2 and ACE2 in comparison with the interactions between RBD1 and ACE2 with Y455, L486 and T501 residues on RBD1. Note that L486F, Y498Q and D501 (instead of T501) to N mutations were found when the sequences of bat coronavirus RaTG13 and SARS-CoV-2 were compared.^2^ Other key residues of RBD2 identified previously,^2^ are Q493 and S494 of which Q493 is also seen to have stable interactions with ACE2 (**Figure 2b** and Supplementary **Figure S2**), while S494 was not detectable in both RBD1-ACE2 and RBD2-ACE2 interactions, suggesting that S494 may not play any critical role at the interface as proposed in the previous study. New residues such as K417, Y473 and A475, which were not reported earlier^2^ but proposed to be important in others work^5,8,13^, emerged to be important for RBD2-ACE2 interface and enhanced the interaction compared to RBD1-ACE2 with V417 and P475 residues. Particularly, the V417K mutation created a salt-bridge with D30 (on ACE2), thus contributing to an increase in the electrostatic interactions^8^ and compensating for coulomb energy due to mutation R439N and loss of distal R439-E329 electrostatic interaction (seen in RBD1-ACE2 interface).^13^ Overall, regardless of the FFs there are the same key clusters of residues in RBD2-ACE2 complex responsible for enhancing the interactions at the interface of RBD2-ACE2 complex; however, the AMBER-FF model yielded a measurable stronger interface with higher number of stable pairs than the C36-FF model.

### Energetics of RBD2-ACE2 vs RBD1-ACE2 interfaces

To energetically differentiate the interfaces, we computed the probability *ρ*(*R*_RBD-ACE2_) for a distance *R*_RBD-ACE2_ between the center of mass of an RBD and ACE2; and a free energy was computed as –*k*_B_*T*log(*ρ*(*R*_RBD-ACE2_)), where *k*_B_ is the Boltzmann constant and *T* = 310 K is the simulated temperature. **Figure 3** shows the free-energy profiles computed for the RBD1-ACE2 and RBD2-ACE2 complexes using the two FFs. When modeled with the C36 FF, the global free-energy profile of RBD1-ACE2 complex with a global minimum (at *R*_RBD-ACE2_ = 47-48.5Å) was broader than RBD2-ACE2 complex that showed a global minimum (at *R*_RBD-ACE2_ ≈ 50 Å) (**Figure 3)**. This suggests that the RBD1-ACE2 interface is less tight than the RBD2-ACE2 interface. When modeled by the AMBER FF, the global energy minima of the two complexes are also located at different positions (*R*_RBD-ACE2_ = 44 Å vs. *R*_RBD-ACE2_ = 47 Å); and the shape of the free-energy profile of RBD1-ACE2 complex for *R*_RBD-ACE2_ > 44 Å is approximately linear with *R*, in contrast to the parabolic shape of RBD2-ACE2 complex. In consistency with the C36-FF model, this suggests that, the RBD2-ACE2 interface has stronger interactions than the RBD1-ACE2 interface. By comparing the free-energy profiles of each complex modeled by the two FFs, we observed that the complex modeled by the AMBER FF has stronger interactions than the complex modeled by the C36 FF. For instance, RBD2-ACE2 complex, if moved apart from the global minimum by 2 Å, shows a free energy increase of 4 *k*_B_*T* in the AMBER-FF model, but only about 2 *k*_B_*T* in the C36-FF model (**Figure 3a**). This is consistent with the above observation based on the pairs of residues at the interfaces (**Figure 2b** and Supplementary **Figure S2**). To this end, even though we observed distinguishable features of the interactions modeled by the two FFs, one similarity stands out clear: the RBD2-ACE2 interface has stronger interaction energies than the RBD1-ACE2 interface as concluded in previous studies,^2–5,8,10–13,30^ which results in more stable pairs of interacting residues in the RBD2-ACE2 interface than the RBD1-ACE2 interface as observed in **Figure 2**.

**Figure 3.**
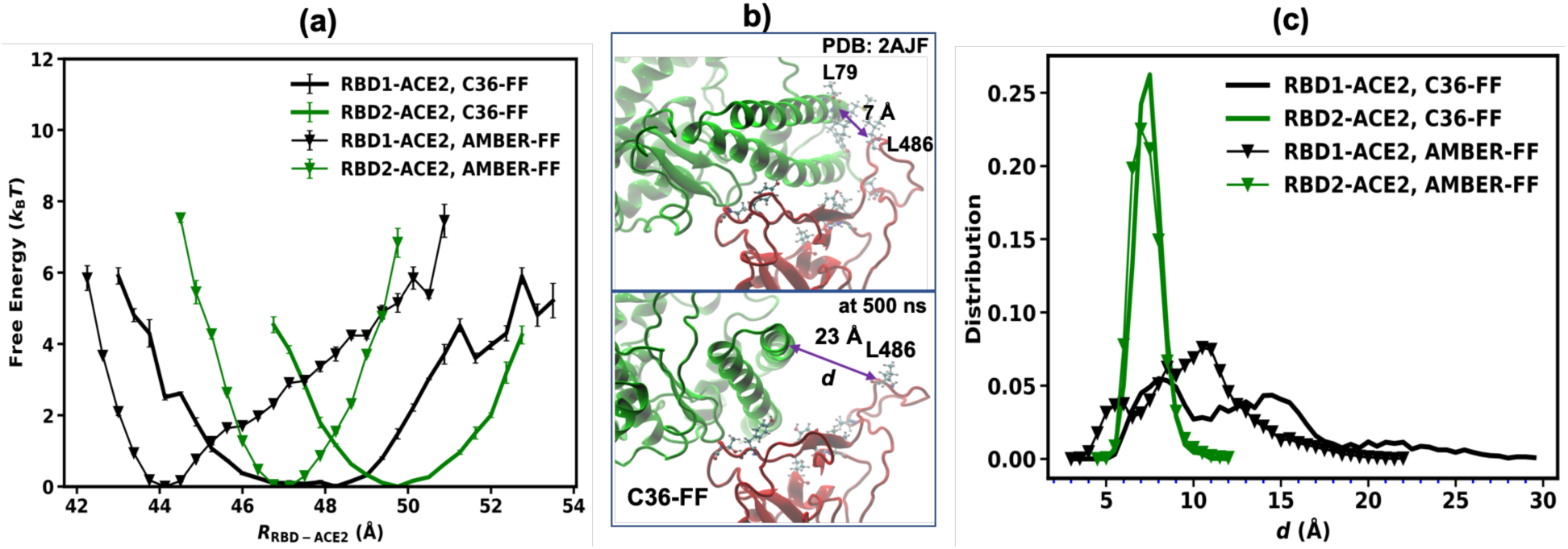
(a) Free-energy profiles computed as function of the distance between the centers of mass of the RBDs and ACE2. (b) Snap-shots showing an initial configuration of RBD1-ACE complex and its configuration at 500 ns from the simulations using C36 FF. This 500 ns configuration was reproducible and showed in 3 out of 8 replicates (c) Distribution of the distance *d* between the Cα of residue L486 of RBD1 and residue L79 of ACE2 and compared with F486-L79 interaction in RBD2-ACE2.

Particularly for RBD1-ACE2 complex, both FFs apparently yield a meta-stable state, which is not evident in the RBD2-ACE2 complex. To the best of our knowledge, this observation has not been reported in any previous studies. To partially explain such meta-stable state observed in the C36-FF, **Figure 3b** shows a snapshot compared to a dynamical structure of RBD1 and its X-ray structure: initially, the loop having L486 and N487 is in close contact to the residues (L79, M82, Y83) of ACE2; at 0.5 μs, it loses its grip on ACE2 and moves up to 23 Å away from ACE2 residues. This is not observed in case of RBD2-ACE2 complex, which remains similar to the crystal structure as shown in **Figure 1b**. To further quantify the behavior of this loop, we measured the distance *d* between the Cα atoms of L486 in RBD1 (or F486 in RBD2) and L79 of ACE2 during the course of simulations. **Figure 3c** shows that in case of RBD1-ACE2 complex, while there is the highest peak in the distribution obtained from the AMBER-FF at *d =* 11 Å, the broader distribution obtained from the C36-FF has two peaks at *d =* 8 Å and *d =* 15 Å, and non-zero values at *d* ∼25 Å. This is also consistent with the above observations that the C36-FF models weaker interactions, particularly between the RBD1 and ACE2 at *d* ∼25 Å than the AMBER-FF. In RBD2-ACE2 complex, the distance is very stable at 7-7.5 Å as indicated by the very sharp peak in simulations using both FFs (**Figure 3c**). However, in RBD1-ACE2 complex, the distributions shift to larger distances in case of both FFs. This suggests that the localized F486 at RBD2-ACE2 interface may have a significantly stronger grip on ACE2 than the corresponding non-polar residue L486 in RBD1.

To this end, these results imply that the mutation L486F in RBD2 of SARS-CoV-2 might be the most consequential as F486 on SARS-CoV-2 has a more stable interaction with the ACE2 helices than L486 on SARS-CoV (**Figure 3b**); the L486 in RBD1 can move away from ACE2, resulting into less effective binding.

### Potential of Mean Force (PMF)

In order to shed more insight into the dissociation events of the RBDs from ACE2, we computed the PMFs of the two complexes via umbrella sampling (US) simulations using the C36 FF (see Methods). Note that the free-energy profiles shown in **Figure 3a** can be poorly sampled at distances away from the global minima, while the US sampling helps to improve rare statistics at those distances.^26–28,31,32^ **Figure 4a** shows that the PMFs for completely dissociating the RBDs from ACE2 around *R*_RBD-ACE2_ = 70 Å are markedly different (the finite size effects^33^ were not excluded from the profiles; so, there is a slight increase at the end of the curves due to the periodic images). We also observed that the meta-stable states in the RBD1-ACE2 complex (**Figure 3a**) become much shallower at *R* = 52 Å (**Figure 4a)**. The dissociation free energy, Δ*G* can be measured as the free-energy difference between *R* = 70 Å and *R* ∼ 50 Å, which is the global free-energy minimum. Our calculation of Δ*G* shows that it costs Δ*G*_1_ = 4.3 ± 0.8 and Δ*G*_2_ = 15.0 ± 0.8 kcal/mol for RBD1 and RBD2 respectively to completely dissociate from ACE2. In other words, the binding of RBD2 to ACE2 yields about 11 kcal/mol more energy than the binding of RBD1. This ΔΔ*G*_MD_ (= Δ*G*_2_ – Δ*G*_1_) is almost three times the value estimated by a coarse-grain model (4.3 kcal/mol) in a recently published work.^12^ This discrepancy may largely come from the simulated parameters (or FFs). It should be noted that ΔΔ*G*_MD_ was computed without a thorough sampling of relative rotational and conformational dynamics of the RBDs and ACE2. The calculations also assumed that the entire conformations of both RBDs and ACE2 are unchanged before the binding. These dynamics if accounted may change the absolute value of ΔΔ*G*_MD_, but likely ΔΔ*G*_MD_ remains positive, indicating that RBD2-ACE2 complex is tighter than RBD1-ACE2 complex.

**Figure 4.**
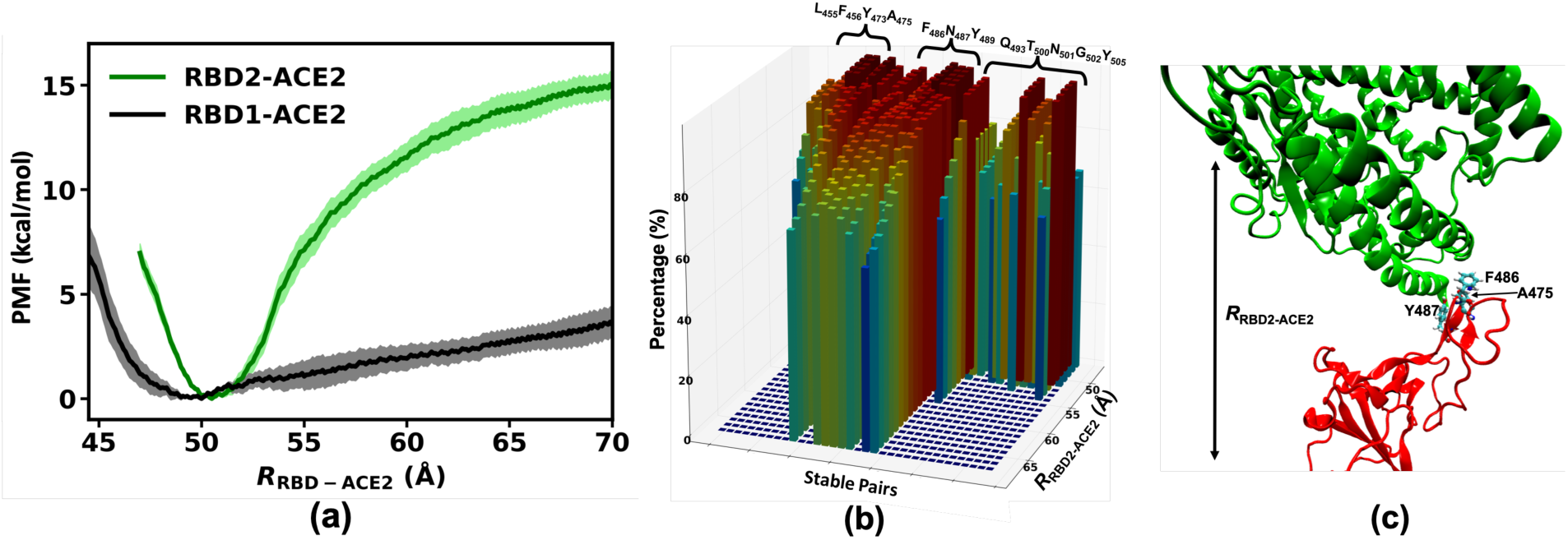
(a) Free-energy profiles computed from US simulations as function of *R*_RBD2-ACE2_ (see Methods). (b) Three-dimension distributions of the stable pairs (see **Figure 2**) as function of the biasing distance between the centers of mass of RBD2 and ACE2. (c) A snapshot during the biasing simulations using *R*_RBD2-ACE2_ = 70 Å. Residue A475 in RBD2 is located right behind F486.

In order to determine which residues of RBD2 are critical and behave collectively during the dissociation of the complex, we computed the probabilities of the stable residue pairs (**Figure 2**) at the interface as a function of the biasing distance between the two centers of mass of RBD2 and ACE2 (**Figure 4b**). **Figure 4b** shows that while the two clusters of residues (L455/F456/Y473 and Q493/T500/N501/G502/Y505) of RBD2 dissociate from ACE2 at *R* = 55 Å (5 Å away from the global minimum), the cluster of residues F486/N487/Y489/A475 (**Figure 4c**) remains in contact with ACE2 with a probability of 60% at *R* = 70 Å. Note that the similar pattern of the changes in the probabilities of these residues indicate that, this cluster of residues behaves as a “sticky” anchor of the Spike protein of SARS-CoV-2 that holds on the binding region of ACE2. In contrast, as observed in **Figure 3b-c**, the similar cluster of L486/N487/Y489/P475 in RBD1 does not have a strong grip on ACE2. These clearly distinguishable outcomes suggest that the mutations L486F and P475A play a critical role that makes RBD2 noticeably stickier to ACE2 than RBD1. In other words, the dynamics of these key residues add a new mechanistic picture of why SARS-CoV-2 is more infectious than SARS-CoV.

## CONCLUSIONS

In this study, we present a simulation data of approximately 20 μs in total (2×5 μs equilibration and 2×4.8 μs US simulations), which show key stable pairs between the SARS-CoV or SARS-CoV-2 RBDs (**Figure 2**) and the human receptor ACE2. We showed three clusters of residues that have enhanced interactions in RBD2 and ACE2. Particularly, one cluster of residues, including F486/N487/Y489/A475, is perhaps the most critical cluster of amino acids that appear to collectively induce strong interactions between RBD2 and ACE2 (**Figure 3c** and **Figure 4**). This cluster of residues could act as an anchor that strongly holds on the N-terminal part of ACE2. In contrast, the corresponding set of residues L486/N487/Y489/P475 of RBD1 can loosen its grip from ACE2 (**Figure 3b-c**). The observations were consistent across different forcefields. The dramatic difference in the dynamics of these residue clusters is only observed in MD simulations of beyond 500 ns, which most previous studies^2–5,8,10–13,30^ did not achieve. The results build our understanding of key determinants of molecular recognition of SARS-CoV-2 and human receptor protein and brings opportunities to target the residue clusters for specific and sensitive binders for therapeutics and diagnostics alike.

## ASSOCIATED CONTENT

### Supporting Information

Figure S1: Distributions of *R*_RBD-ACE2_ from the US sampling simulations. Figure S2. Pairs of residues at the RBD1-ACE2 and RBD2-ACE2 interfaces modeled by the AMBER ff88SB FF

## ACKNOWLEDGMENT

V.A.N is a Director’s Postdoctoral Fellow at LANL and is partially funded by Laboratory Directed R&D Postdoctoral Research and Development fellowship (20170692PRD4). The work was authored under Triad National Security, LLC (“Triad”) Contract No. 89233218CNA000001 with the U.S. Department of Energy. This research used computational resources provided by the Los Alamos National Laboratory Institutional Computing Program (under w20_foldamers to R.K.J. and x20_simcovid19 to V.A.N.), which is supported by the U.S. Department of Energy National Nuclear Security Administration under Contract No. 89233218CNA000001.

## ABBREVIATIONS

ACE2: angiotensin-converting enzyme 2
covid: coronavirus disease
MD: molecular dynamics
PMF: potential of mean force
RBD: receptor binding domain
RBM: receptor binding motif
SARS: severe acute respiratory syndrome
FF: forcefield
US: umbrella sampling

## For Table of Contents Use Only

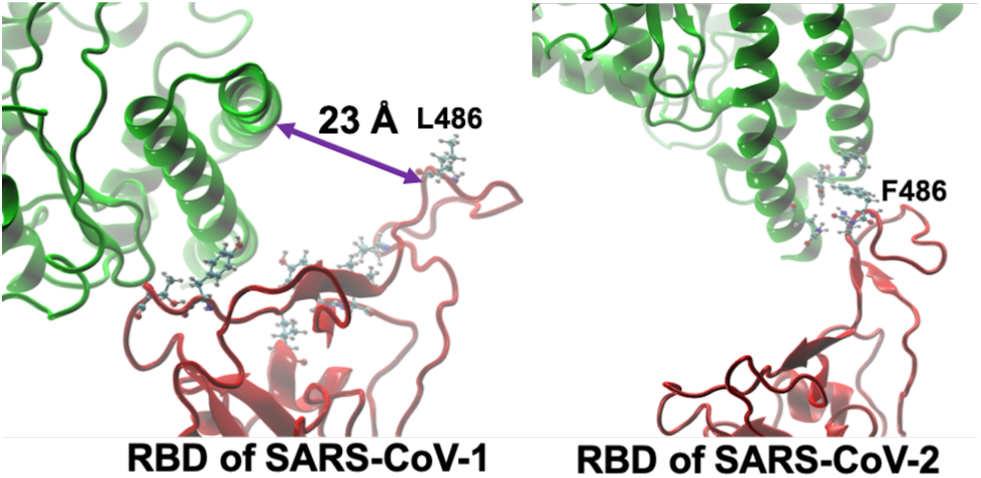

## Supporting Information for

**Figure S1.**
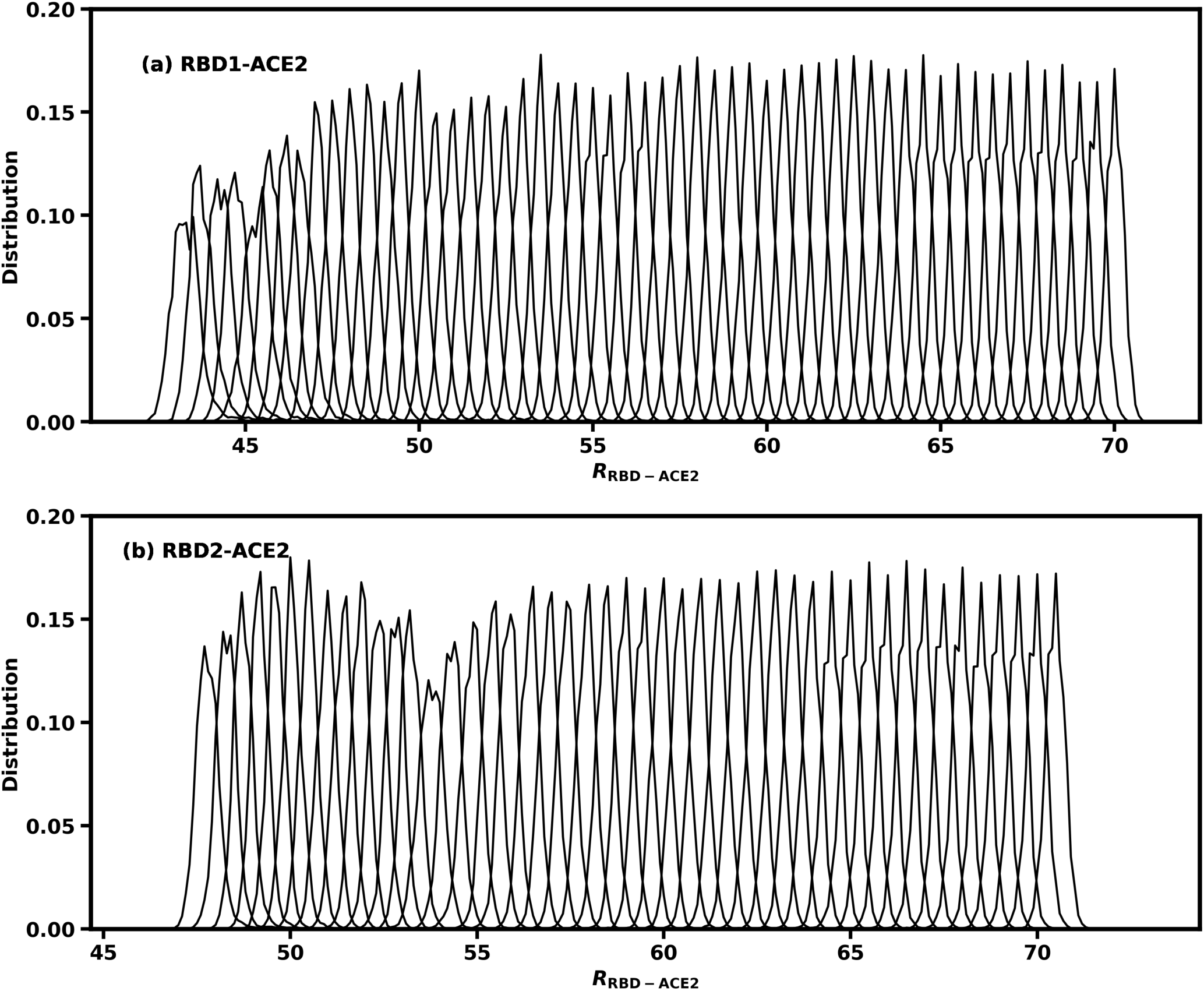
Distribution of *R*_RBD-ACE2_ computed from the US sampling.

**Figure S2.**
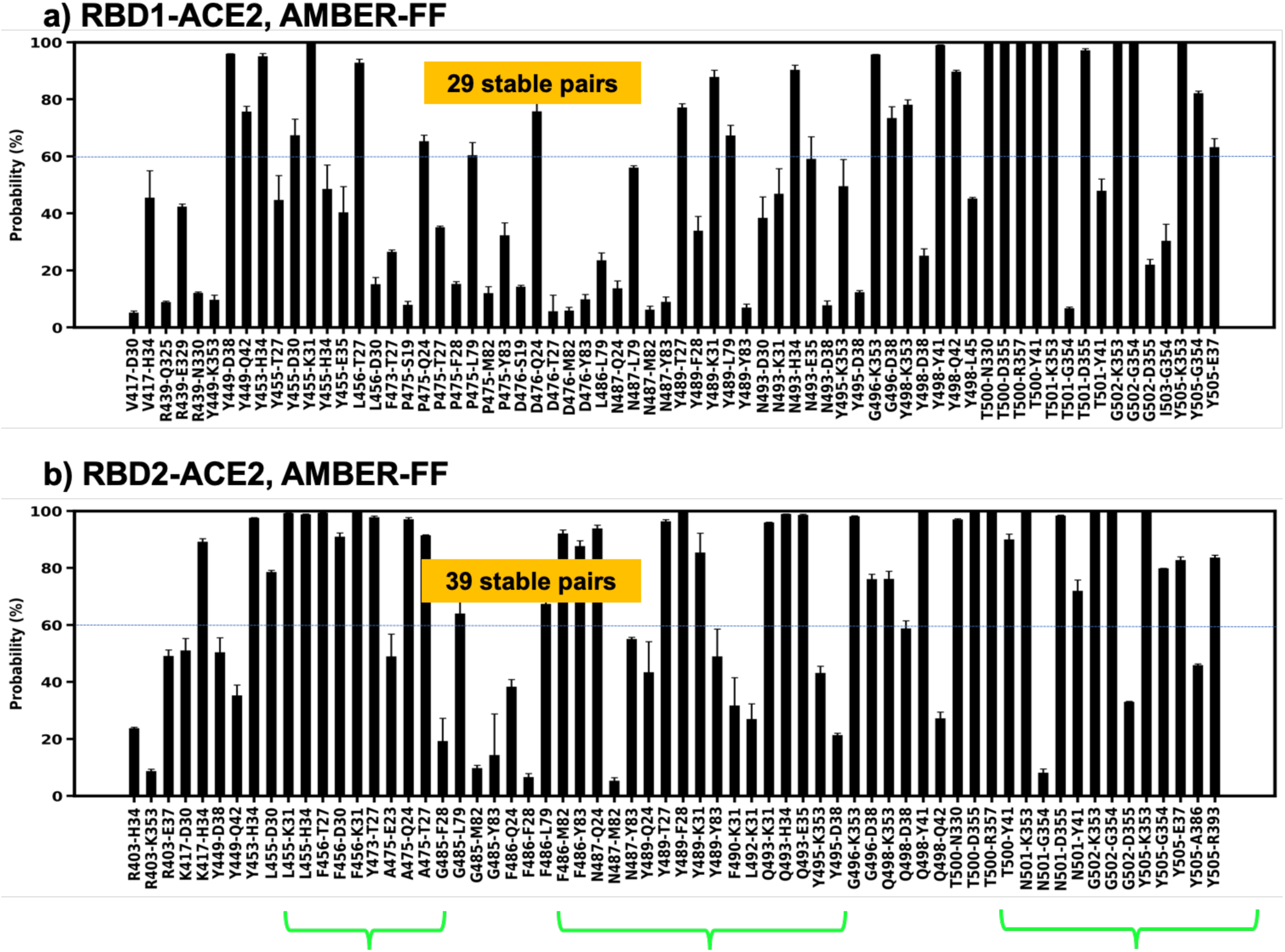
(a-b) Probability of pairs of residues found within 3.5 Å at the interfaces between RBD1-2 with ACE2 during simulations of 10 μs per system using AMBER ff88SB FF^15^ performed by the Shaw group.^7^ A stable pair is defined to have a probability larger than 60% during the simulation times, i.e, forming a pair within 3.5 Å for more than 60% of a simulation trajectory. Green brackets indicate clusters of residues that are on average more stable in RBD2-ACE2 interface than the corresponding clusters in RBD1-ACE2 interface.

